## Supporting information File 1. Clavis JSON-schema for "Clavis: an open and versatile identification key format"

```
{
  "$schema": "http://json-schema.org/draft-07/schema#",
  "title": "Clavis identification key schema",
  "description": "Clavis-compliant keys contain knowledge that may be
used to distinguish taxa from each other.",
  "type": "object",
  "required": [
    "$schema",
    "title",
    "language",
    "license",
    "creator",
    "lastModified",
    "identifier",
    "taxa",
    "characters",
    "statements",
    "persons"
  ],
  "properties": {
    "$schema": {
      "description": "The schema url of (this) schema defining the
format of the key.",
      "$ref": "#/definitions/url"
    },
    "title": {
      "description": "The name of the key",
      "comment": "Accepts array for multilingual support.",
      "$ref": "#/definitions/localizedString",
      "examples": [
        "Birds of Norway"
      ]
    },
    "media": {
      "description": "The logo/illustration image of the key.",
      "$ref": "#/definitions/mediaID"
    },
    "description": {
      "description": "Short description of the key (valid markdown).",
      "comment": "Accepts array for multilingual support.",
      "$ref": "#/definitions/localizedString",
      "contentMediaType": "text/markdown"
    },
    "descriptionDetails": {
      "description": "Extended description of the key that supplements
the description (valid markdown).",
```

```
      "comment": "Accepts array for multilingual support.",
      "$ref": "#/definitions/localizedString",
      "contentMediaType": "text/markdown"
    },
    "descriptionUrl": {
      "description": "Hyperlink to more information on the key (valid
url).",
      "comment": "Accepts array for multilingual support.",
      "$ref": "#/definitions/localizedUrl"
    },
    "audience": {
      "description": "Description of the intended audience for the
key.",
      "comment": "Accepts array for multilingual support.",
      "$ref": "#/definitions/localizedString",
      "examples": [
        "Undergraduate students and up."
      ]
    },
    "source": {
      "description": "Source of the key.",
      "comment": "Accepts array for multilingual support.",
      "$ref": "#/definitions/localizedString",
      "examples": [
        "Koch, Wouter (2019). Birds of Norway. ISBN 1234567890"
      ]
    },
    "sourceUrl": {
      "description": "Hyperlink to the source of the key (valid
url).",
      "comment": "Accepts array for multilingual support.",
      "$ref": "#/definitions/localizedUrl",
      "examples": [
        "https://doi.org/10.1126/science.1251554"
      ]
    },
    "geography": {
      "description": "The region for which the key is valid (e.g.
covers all subtaxa), represented as a geography object.",
      "$ref": "#/definitions/geography"
    },
    "primaryContact": {
      "description": "The organization- or person-id that is the main
contact point for the key.",
      "oneOf": [
        {
          "$ref": "#/definitions/personID"
        },
        {
          "$ref": "#/definitions/organizationID"
        }
      ]
```

```
    },
    "creator": {
      "description": "The id(s) of the creator(s) of the key",
      "oneOf": [
        {
          "$ref": "#/definitions/personID"
        },
        {
          "type": "array",
          "items": {
            "$ref": "#/definitions/personID"
          }
        }
      ]
    },
    "contributor": {
      "description": "The id(s) of the contributor(s) of the key",
      "oneOf": [
        {
          "$ref": "#/definitions/personID"
        },
        {
          "type": "array",
          "items": {
            "$ref": "#/definitions/personID"
          }
        }
      ]
    },
    "publisher": {
      "description": "The id(s) of the publishing institutions of the
key.",
      "oneOf": [
        {
          "$ref": "#/definitions/organizationID"
        },
        {
          "type": "array",
          "items": {
            "$ref": "#/definitions/organizationID"
          }
        }
      ]
    },
    "license": {
      "description": "The url to the license under which the key
falls.",
      "$ref": "#/definitions/url",
      "examples": [
        "https://creativecommons.org/licenses/by/4.0/"
      ]
    },
```

```
    "language": {
      "description": "The ISO 639-1 code(s) of the key language(s).",
      "comment": "String for a single language, array of strings for multilingual support. If used as an array, be sure to use the localizedString and localizedUrl as arrays too.",
      "oneOf": [
        {
          "type": "string",
          "pattern": "^[a-z]{2}$"
        },
        {
          "type": "array",
          "items": {
            "type": "string",
            "pattern": "^[a-z]{2}$"
          }
        }
      ],
      "examples": [
        "en",
        "nb",
        [
          "en",
          "nb"
        ]
      ]
    },
    "created": {
      "description": "The moment the key was made or first published, as 'YYYY-MM-DD hh:mm:ss'.",
      "type": "string",
      "pattern": "^20\\d\\d-(0[1-9]|1[0-2])-([012]\\d|3[01]) ([01]\\d|2[0-3]):([0-5]\\d):([0-5]\\d)$",
      "examples": [
        "2019-05-21 22:51:55"
      ]
    },
    "lastModified": {
      "description": "The most recent moment the key was modified, as 'YYYY-MM-DD hh:mm:ss'.",
      "type": "string",
      "pattern": "^20\\d\\d-(0[1-9]|1[0-2])-([012]\\d|3[01]) ([01]\\d|2[0-3]):([0-5]\\d):([0-5]\\d)$",
      "examples": [
        "2019-05-21 22:51:55"
      ]
    },
    "identifier": {
      "description": "The GUID of this key (persistent regardless of version).",
      "type": "string"
    },
```

```
    "url": {
      "description": "The url of where the key lives (to check for
newer versions).",
      "$ref": "#/definitions/url"
    },
    "externalServices": {
      "description": "Services used by the key for lookups of images,
taxa, etc.",
      "type": "array",
      "items": {
        "$ref": "#/definitions/externalService"
      }
    },
    "userRequirements": {
      "description": "Requirements to the users of the various
characters, so that the user can be warned, helped, etc.",
      "type": "array",
      "items": {
        "$ref": "#/definitions/userRequirement"
      }
    },
    "taxa": {
      "description": "Taxa (e.g. species) the key can resolve to. Do
not have to be exclusively taxonomic units.",
      "comment": "Taxa to which the key can resolve (either the taxa
directly or their children).",
      "type": "array",
      "items": {
        "$ref": "#/definitions/taxon"
      }
    },
    "characters": {
      "description": "Characters (questions, e.g. 'Wing color' or
'Number of spots') used to distinguish between two or more taxa.",
      "type": "array",
      "items": {
        "$ref": "#/definitions/character"
      }
    },
    "statements": {
      "description": "Relationships between taxa and character states
(or lack thereof) that define those taxa.",
      "type": "array",
      "items": {
        "$ref": "#/definitions/statement"
      }
    },
    "persons": {
      "description": "Persons that are connected to (parts of) the
key, such as creators.",
      "type": "array",
      "items": {
```

```
          "$ref": "#/definitions/person"
        }
      },
      "organizations": {
        "description": "Organizations that are connected to (parts of)
the key or persons, such as employers and publishers.",
        "type": "array",
        "items": {
          "$ref": "#/definitions/organization"
        }
      },
      "mediaElements": {
        "description": "Media elements that are used in the key.",
        "type": "array",
        "items": {
          "$ref": "#/definitions/localizedMediaElement"
        }
      }
    },
    "additionalProperties": false,
    "definitions": {
      "localizedString": {
        "description": "Language-dependent string or object of strings,
with keys corresponding to the languages supported by the key.",
        "oneOf": [
          {
            "type": "string"
          },
          {
            "type": "object",
            "propertyNames": {
              "pattern": "^[a-z]{2}$"
            },
            "properties": {},
            "additionalProperties": {
              "type": "string"
            }
          }
        ]
      },
      "localizedUrl": {
        "description": "Language-dependent urls or object of urls,
corresponding to the languages supported by the key.",
        "oneOf": [
          {
            "$ref": "#/definitions/url"
          },
          {
            "type": "object",
            "propertyNames": {
              "pattern": "^[a-z]{2}$"
            },
```

```
          "properties": {},
          "additionalProperties": {
            "$ref": "#/definitions/url"
          }
        }
      ]
    },
    "localizedMediaElement": {
      "type": "object",
      "description": "Language-dependent media element or object of
media elements, corresponding to the languages supported by the key.",
      "properties": {
        "id": {
          "description": "Internally unique id of the localized media
element.",
          "$ref": "#/definitions/mediaID"
        },
        "mediaElement": {
          "description": "The media element or media elements (one for
each language).",
          "oneOf": [
            {
              "$ref": "#/definitions/mediaElement"
            },
            {
              "type": "object",
              "propertyNames": {
                "pattern": "^[a-z]{2}$"
              },
              "properties": {},
              "additionalProperties": {
                "$ref": "#/definitions/mediaElement"
              }
            }
          ]
        }
      },
      "additionalProperties": false
    },
    "url": {
      "description": "String formed as a url, or an external
resource.",
      "oneOf": [
        {
          "type": "string",
          "format": "uri"
        },
        {
          "$ref": "#/definitions/externalResource"
        }
      ]
    },
```

```
    "taxonID": {
      "description": "String used as an internal ID for a taxon.
Lowercase alphanumeric and underscores are allowed.",
      "type": "string",
      "pattern": "^taxon:[a-z0-9_]+$"
    },
    "characterID": {
      "description": "String used as an internal ID for a character.
Lowercase alphanumeric and underscores are allowed.",
      "type": "string",
      "pattern": "^character:[a-z0-9_]+$"
    },
    "stateID": {
      "description": "String used as an internal ID for a state.
Lowercase alphanumeric and underscores are allowed.",
      "type": "string",
      "pattern": "^state:[a-z0-9_]+$"
    },
    "personID": {
      "description": "String used as an internal ID for a person.
Lowercase alphanumeric and underscores are allowed.",
      "type": "string",
      "pattern": "^person:[a-z0-9_]+$"
    },
    "organizationID": {
      "description": "String used as an internal ID for an
organization. Lowercase alphanumeric and underscores are allowed.",
      "type": "string",
      "pattern": "^organization:[a-z0-9_]+$"
    },
    "serviceID": {
      "description": "String used as an internal ID for a service.
Lowercase alphanumeric and underscores are allowed.",
      "type": "string",
      "pattern": "^service:[a-z0-9_]+$"
    },
    "statementID": {
      "description": "String used as an internal ID for a statement.
Lowercase alphanumeric and underscores are allowed.",
      "type": "string",
      "pattern": "^statement:[a-z0-9_]+$"
    },
    "userRequirementID": {
      "description": "String used as an internal ID for a user
requirement. Lowercase alphanumeric and underscores are allowed.",
      "type": "string",
      "pattern": "^requirement:[a-z0-9_]+$"
    },
    "mediaID": {
      "description": "String used as an internal ID for a media
element. Lowercase alphanumeric and underscores are allowed.",
      "type": "string",
```

```
      "pattern": "^media:[a-z0-9_]+$"
    },
    "mediaFile": {
      "type": "object",
      "properties": {
        "title": {
          "description": "The title of the media file.",
          "$ref": "#/definitions/localizedString"
        },
        "url": {
          "description": "The reference to the media file (url or
resource).",
          "$ref": "#/definitions/url"
        },
        "file": {
          "description": "The actual media file (base64 or svg) as a
data URI scheme.",
          "oneOf": [
            {
              "type": "string",
              "pattern":
"^data:([a-z0-9/]+);base64,([a-zA-Z0-9+/=]+)$"
            },
            {
              "type": "string",
              "pattern": "^data:image/svg\\+xml;utf8,(.*)$"
            }
          ]
        },
        "width": {
          "description": "The number of pixels horizontally (if a
bitmap image or video).",
          "type": "integer"
        },
        "height": {
          "description": "The number of pixels vertically (if a bitmap
image or video).",
          "type": "integer"
        },
        "length": {
          "description": "The length in seconds of an audio or video
file.",
          "type": "integer"
        },
        "placeholder": {
          "description": "Image file that can be shown instead of the
video or audio file.",
          "$ref": "#/definitions/mediaID"
        },
        "creator": {
          "description": "The id(s) of the creator(s) of the media
file",
```

```
     "oneOf": [
      {
       "$ref": "#/definitions/personID"
      },
      {
       "type": "array",
       "items": {
        "$ref": "#/definitions/personID"
       }
      }
     ]
    },
    "contributor": {
     "description": "The id(s) of the contributor(s) of the media
file",
     "oneOf": [
      {
       "$ref": "#/definitions/personID"
      },
      {
       "type": "array",
       "items": {
        "$ref": "#/definitions/personID"
       }
      }
     ]
    },
    "publisher": {
     "description": "The id(s) of the publishing institutions of
the media file.",
     "oneOf": [
      {
       "$ref": "#/definitions/organizationID"
      },
      {
       "type": "array",
       "items": {
        "$ref": "#/definitions/organizationID"
       }
      }
     ]
    },
    "license": {
     "description": "The url to the license under which the media
file falls.",
     "$ref": "#/definitions/url",
     "examples": [
      "https://creativecommons.org/licenses/by/4.0/"
     ]
    }
   },
   "additionalProperties": false
```

```
  },
  "mediaElement": {
   "description": "A media element (collection of various formats
of the same media object).",
   "type": "object",
   "properties": {
    "file": {
     "description": "The various formats of the same media
object.",
     "oneOf": [
      {
       "$ref": "#/definitions/mediaFile"
      },
      {
       "type": "array",
       "items": {
        "$ref": "#/definitions/mediaFile"
       }
      }
     ]
    }
   },
   "additionalProperties": false
  },
  "multiPolygon": {
   "description": "The coordinates array of a GeoJSON
MultiPolygon.",
   "type": "array",
   "items": {
    "type": "array",
    "items": {
     "type": "array",
     "items": {
      "type": "array",
      "items": {
       "type": "number"
      }
     }
    }
   }
  },
  "geography": {
   "description": "A geographic element (name, polygon, and/or
external service).",
   "type": "object",
   "properties": {
    "name": {
     "description": "The name of the area(s).",
     "comment": "Accepts array for multilingual support.",
     "$ref": "#/definitions/localizedString",
     "examples": [
      "Norway",
```

```
            "Europe",
            "Trøndelag",
            [
              "Norge",
              "Norway"
            ]
          ]
        },
        "polygon": {
          "description": "The geographical area(s), represented as the
coordinates array of a GeoJSON MultiPolygon.",
          "$ref": "#/definitions/multiPolygon"
        },
        "service": {
          "description": "An url or external service that returns
geographical information.",
          "$ref": "#/definitions/url"
        }
      },
      "additionalProperties": false
    },
    "externalResource": {
      "description": "A resource managed elsewhere.",
      "type": "object",
      "properties": {
        "serviceId": {
          "description": "The id to one of the externalServices
defined.",
          "$ref": "#/definitions/serviceID"
        },
        "externalId": {
          "description": "The id of the resource at the
externalService.",
          "type": "string"
        }
      },
      "additionalProperties": false
    },
    "externalService": {
      "description": "Service used by the key, for media files,
taxonomy and/or nomenclature, species distributions, etc.",
      "type": "object",
      "required": [
        "id"
      ],
      "properties": {
        "id": {
          "description": "Internally unique id to the service.",
          "$ref": "#/definitions/serviceID"
        },
        "title": {
          "description": "Name of the service.",
```

```
          "type": "string"
        },
        "description": {
          "description": "Description of the service.",
          "type": "string"
        },
        "provider": {
          "description": "Provider of the service.",
          "type": "string"
        },
        "url": {
          "description": "Url for the service documentation.",
          "$ref": "#/definitions/url"
        }
      },
      "additionalProperties": false
    },
    "person": {
      "type": "object",
      "required": [
        "id",
        "name"
      ],
      "properties": {
        "id": {
          "$ref": "#/definitions/personID"
        },
        "name": {
          "description": "Full name of the person",
          "comment": "Accepts object for multilingual support.",
          "$ref": "#/definitions/localizedString"
        },
        "email": {
          "description": "Email address of the person",
          "type": "string",
          "format": "email"
        },
        "url": {
          "description": "Hyperlink to more information on the person
(valid url).",
          "comment": "Accepts object for multilingual support.",
          "$ref": "#/definitions/localizedUrl"
        },
        "media": {
          "description": "A media file (image) representing the
person.",
          "$ref": "#/definitions/mediaID"
        },
        "affiliation": {
          "description": "Organization id(s) the person is affiliated
with.",
          "oneOf": [
```

```
            {
              "$ref": "#/definitions/organizationID"
            },
            {
              "type": "array",
              "items": {
                "$ref": "#/definitions/organizationID"
              }
            }
          ]
        }
      },
      "additionalProperties": false
    },
    "organization": {
      "type": "object",
      "required": [
        "id",
        "name"
      ],
      "properties": {
        "id": {
          "$ref": "#/definitions/organizationID"
        },
        "name": {
          "description": "Name of the organization",
          "comment": "Accepts object for multilingual support.",
          "$ref": "#/definitions/localizedString"
        },
        "url": {
          "description": "Hyperlink to more information on the
organization (valid url).",
          "comment": "Accepts object for multilingual support.",
          "$ref": "#/definitions/localizedUrl"
        },
        "primaryContact": {
          "description": "The person-id that is the main contact point
for the organization.",
          "$ref": "#/definitions/personID"
        },
        "media": {
          "description": "A media file (image) representing the
organization, such as a logo.",
          "$ref": "#/definitions/mediaID"
        }
      },
      "additionalProperties": false
    },
    "userRequirement": {
      "type": "object",
      "required": [
        "id"
```

```
      ],
      "properties": {
        "id": {
          "$ref": "#/definitions/userRequirementID"
        },
        "title": {
          "comment": "Accepts array for multilingual support.",
          "$ref": "#/definitions/localizedString"
        },
        "warning": {
          "comment": "Accepts array for multilingual support.",
          "$ref": "#/definitions/localizedString"
        },
        "description": {
          "description": "Short description of the requirements to the
user (valid markdown).",
          "comment": "Accepts object for multilingual support.",
          "$ref": "#/definitions/localizedString",
          "contentMediaType": "text/markdown"
        },
        "descriptionDetails": {
          "description": "Extended description of the requirements to
the user that supplements the description (valid markdown).",
          "comment": "Accepts object for multilingual support.",
          "$ref": "#/definitions/localizedString",
          "contentMediaType": "text/markdown"
        },
        "descriptionUrl": {
          "description": "Hyperlink to more information on the
requirements to the user (valid url).",
          "comment": "Accepts object for multilingual support.",
          "$ref": "#/definitions/localizedUrl"
        },
        "media": {
          "description": "Media or illustration that informs the user
on the requirements to the user.",
          "comment": "Accepts object for multilingual support.",
          "$ref": "#/definitions/mediaID"
        }
      },
      "additionalProperties": false
    },
    "taxon": {
      "type": "object",
      "oneOf": [
        {
          "required": [
            "id",
            "scientificName"
          ]
        },
        {
```

```
          "required": [
            "id",
            "externalReference"
          ]
        },
        {
          "required": [
            "id",
            "label"
          ]
        }
      ],
      "properties": {
        "id": {
          "description": "Internally unique id to the taxon.",
          "$ref": "#/definitions/taxonID"
        },
        "scientificName": {
          "description": "Scientific name of the taxon.",
          "minLength": 5,
          "type": "string",
          "examples": [
            "Vulpes lagopus"
          ]
        },
        "scientificNameAuthor": {
          "description": "Author string of the scientific name of the
taxon.",
          "type": "string",
          "examples": [
            "Koch, 1888"
          ]
        },
        "placeholderName": {
          "description": "Name that can be shown while fetching the
name externally. Also useful for editing the key.",
          "comment": "Accepts object for multilingual support.",
          "$ref": "#/definitions/localizedString",
          "examples": [
            "B. hortorum (melanistic queen)"
          ]
        },
        "vernacularName": {
          "description": "Vernacular name of the taxon.",
          "comment": "Accepts object for multilingual support.",
          "$ref": "#/definitions/localizedString",
          "examples": [
            "fjellrev",
            {
              "no": "fjellrev",
              "en": "Arctic Fox"
            }
```

```
          ]
        },
        "media": {
          "description": "Media elements of the taxon.",
          "oneOf": [
            {
              "$ref": "#/definitions/mediaID"
            },
            {
              "type": "array",
              "items": {
                "$ref": "#/definitions/mediaID"
              }
            }
          ]
        },
        "description": {
          "description": "Short description of the taxon (valid
markdown).",
          "comment": "Accepts object for multilingual support.",
          "$ref": "#/definitions/localizedString",
          "contentMediaType": "text/markdown"
        },
        "descriptionDetails": {
          "description": "Extended description of the taxon that
supplements the description (valid markdown).",
          "comment": "Accepts object for multilingual support.",
          "$ref": "#/definitions/localizedString",
          "contentMediaType": "text/markdown"
        },
        "descriptionUrl": {
          "description": "Hyperlink or resource to more information on
the taxon.",
          "comment": "Accepts object for multilingual support.",
          "$ref": "#/definitions/localizedUrl"
        },
        "rank": {
          "description": "Name of the level of the taxon.",
          "comment": "Accepts object for multilingual support.",
          "$ref": "#/definitions/localizedString",
          "examples": [
            "slekt",
            {
              "no": "slekt",
              "en": "genus"
            }
          ]
        },
        "label": {
          "description": "Type of morph of the taxon.",
          "type": "string",
          "minLength": 0,
```

```
     "examples": [
      "male",
      "♀",
      "larva"
     ]
    },
    "isEndPoint": {
     "description": "Whether the key should stop when this taxon
is the only remaining possibility, even when it has multiple children
remaining.",
     "comment": "Default is FALSE (if not specified). A taxon
without children is alway an endpoint by definition, unless one of its
ancestors overrides this by being specified as an endpoint.",
     "type": "boolean"
    },
    "children": {
     "type": "array",
     "items": {
      "$ref": "#/definitions/taxon"
     }
    },
    "externalReference": {
     "description": "Reference to a taxon at one or more
providers, each as an object with a provider id and a taxon id at that
provider.",
     "comment": "Accepts an array for multiple sources. Each
element accepts an object for multilingual support.",
     "oneOf": [
      {
       "$ref": "#/definitions/localizedUrl"
      },
      {
       "type": "array",
       "items": {
        "$ref": "#/definitions/localizedUrl"
       }
      }
     ]
    },
    "followUp": {
     "description": "Url or reference to instance at external
service for a key for this taxon, that for instance can be used to
identify to a lower rank than the current key can.",
     "comment": "Accepts array for multilingual support.",
     "$ref": "#/definitions/localizedUrl"
    },
    "geography": {
     "description": "The area(s) in which the taxon occurs,
represented as a geography object.",
     "$ref": "#/definitions/geography"
    }
   },
```

```
   "additionalProperties": false
  },
  "character": {
   "type": "object",
   "oneOf": [
    {
     "required": [
      "id",
      "title",
      "states"
     ]
    },
    {
     "required": [
      "id",
      "title",
      "type",
      "min",
      "max",
      "stepSize",
      "unit"
     ]
    }
   ],
   "properties": {
    "id": {
     "description": "Internally unique id to the character.",
     "$ref": "#/definitions/characterID"
    },
    "title": {
     "description": "Name of the character.",
     "comment": "Accepts array for multilingual support.",
     "$ref": "#/definitions/localizedString",
     "examples": [
      "Color of the wings"
     ]
    },
    "media": {
     "description": "The media element(s) of the character. Can
be used to inform user of relevant structures etc.",
     "oneOf": [
      {
       "$ref": "#/definitions/mediaID"
      },
      {
       "type": "array",
       "items": {
        "$ref": "#/definitions/mediaID"
       }
      }
     ]
    },
```

```
        "description": {
          "description": "Short description of the character (valid
markdown).",
          "comment": "Accepts object for multilingual support.",
          "$ref": "#/definitions/localizedString",
          "contentMediaType": "text/markdown"
        },
        "descriptionDetails": {
          "description": "Extended description of the character that
supplements the description (valid markdown).",
          "comment": "Accepts object for multilingual support.",
          "$ref": "#/definitions/localizedString",
          "contentMediaType": "text/markdown"
        },
        "descriptionUrl": {
          "description": "Hyperlink or resource to more information on
the character.",
          "comment": "Accepts object for multilingual support.",
          "$ref": "#/definitions/localizedUrl"
        },
        "type": {
          "description": "Type of the character (exclusive when states
are categorical and mutually exclusive, non-exclusive when these are
non-exclusive, or numerical when the state is numerical).",
          "comment": "Default is exclusive (if not specified).",
          "type": "string",
          "enum": [
            "exclusive",
            "non-exclusive",
            "numerical"
          ]
        },
        "userRequirement": {
          "description": "Id to the userRequirement required to answer
this character.",
          "comment": "Has to be one of the userRequirement defined on
the key level.",
          "$ref": "#/definitions/userRequirementID"
        },
        "logicalPremise": {
          "description": "Logical requirement that has to be fulfilled
for this question to be asked.",
          "comment": "Has to refer to stateIds, that have to be fully
true (either answered or all alternatives ruled out). Can use !, &&,
||, (, ), <, >, =.",
          "type": "string",
          "pattern": "^(( && )|( \\|\\|
)|(&&)|(\\|\\|)|[a-z0-9_:()!<>=])+$"
        },
        "min": {
          "type": "number",
          "description": "The minimum numerical value for the
```

```
character."
        },
        "max": {
          "type": "number",
          "description": "The maximum numerical value for the
character."
        },
        "stepSize": {
          "type": "number",
          "description": "The increments with which the numerical
value of the character can be specified."
        },
        "unit": {
          "description": "The unit of the numerical value.",
          "$ref": "#/definitions/localizedString",
          "examples": [
            "mm",
            "meters below the surface",
            "spots",
            "legs",
            "kg"
          ]
        },
        "states": {
          "oneOf": [
            {
              "type": "array",
              "items": {
                "$ref": "#/definitions/state"
              }
            },
            {
              "$ref": "#/definitions/state"
            }
          ]
        }
      },
      "additionalProperties": false
    },
    "state": {
      "description": "The value a character can have.",
      "type": "object",
      "required": [
        "id",
        "title"
      ],
      "properties": {
        "id": {
          "description": "Internally unique id of the state.",
          "$ref": "#/definitions/stateID"
        },
        "title": {
```

```
          "description": "Content of the state.",
          "comment": "Only to be used for categorical characters.
Accepts object for multilingual support.",
          "$ref": "#/definitions/localizedString"
        },
        "media": {
          "description": "Media element(s) that illustrate the
state.",
          "oneOf": [
            {
              "$ref": "#/definitions/mediaID"
            },
            {
              "type": "array",
              "items": {
                "$ref": "#/definitions/mediaID"
              }
            }
          ]
        },
        "description": {
          "description": "Short description of the state (valid
markdown).",
          "comment": "Accepts object for multilingual support.",
          "$ref": "#/definitions/localizedString",
          "contentMediaType": "text/markdown"
        },
        "descriptionDetails": {
          "description": "Extended description of the state that
supplements the description (valid markdown).",
          "comment": "Accepts object for multilingual support.",
          "$ref": "#/definitions/localizedString",
          "contentMediaType": "text/markdown"
        },
        "descriptionUrl": {
          "description": "Hyperlink or resource to more information on
the state.",
          "comment": "Accepts object for multilingual support.",
          "$ref": "#/definitions/localizedUrl"
        }
      },
      "additionalProperties": false
    },
    "statement": {
      "description": "A fact connecting a taxon and a character
through a certain value.",
      "type": "object",
      "required": [
        "id",
        "taxon",
        "character",
        "value",
```

```
        "frequency"
      ],
      "properties": {
        "id": {
          "description": "Internally unique id of the statement.",
          "$ref": "#/definitions/statementID"
        },
        "taxon": {
          "description": "Id of the taxon this statement is about.",
          "$ref": "#/definitions/taxonID"
        },
        "character": {
          "description": "Id of the character this statement is
about.",
          "$ref": "#/definitions/characterID"
        },
        "value": {
          "description": "A value for this character for this taxon.
Must be either the id of a state, or an array of floats [min, max] for
a numerical range.",
          "oneOf": [
            {
              "$ref": "#/definitions/stateID"
            },
            {
              "type": "array",
              "items": {
                "type": "number"
              },
              "minItems": 2,
              "maxItems": 2
            }
          ]
        },
        "frequency": {
          "description": "The frequency with which the taxon has this
value for this character.",
          "type": "number",
          "minimum": 0,
          "maximum": 1
        },
        "geography": {
          "description": "The area(s) in which the taxon can have this
property, represented as a geography object.",
          "$ref": "#/definitions/geography"
        },
        "media": {
          "description": "Illustration(s) of this particular taxon
having this particular property (this value for this character).",
          "oneOf": [
            {
              "$ref": "#/definitions/mediaID"
```

```
            },
            {
              "type": "array",
              "items": {
                "$ref": "#/definitions/mediaID"
              }
            }
          ]
        },
        "description": {
          "description": "Short description of the taxon having this
property (valid markdown).",
          "comment": "Accepts array for multilingual support.",
          "$ref": "#/definitions/localizedString",
          "contentMediaType": "text/markdown"
        },
        "descriptionDetails": {
          "description": "Extended description of the taxon having
this property that supplements the description (valid markdown).",
          "comment": "Accepts array for multilingual support.",
          "$ref": "#/definitions/localizedString",
          "contentMediaType": "text/markdown"
        },
        "descriptionUrl": {
          "description": "Hyperlink or resource to more information on
the taxon having this property.",
          "comment": "Accepts array for multilingual support.",
          "$ref": "#/definitions/localizedUrl"
        }
      },
      "additionalProperties": false
    }
  }
}
```
