## Supplemental Data 1 for "Clavis: an open and versatile identification key format"

### Clavis key example: Pokémon


```
{
  "$schema":
"https://raw.githubusercontent.com/Artsdatabanken/Clavis/main/Schema/C
lavis.json",
  "title": "A key to a selection of Pokémon",
  "language": "en",
  "license": "https://creativecommons.org/licenses/by/4.0/",
  "creator": "person:wouterkoch",
  "lastModified": "2022-10-15 01:31:22",
  "identifier": "26b57071-15ca-4b44-92a4-b61181f15373",
  "persons": [
    {
      "id": "person:wouterkoch",
      "name": "Wouter Koch"
    }
  ],
  "organizations": [
    {
      "id": "organization:ntnu",
      "name": "Norwegian University of Science and Technology",
      "url": "https://www.ntnu.no"
    }
  ],
  "taxa": [
    {
      "id": "taxon:pikachuidae",
      "scientificName": "Pikachuidae",
      "children": [
        {
          "id": "taxon:pokemon_172",
          "scientificName": "Pichu",
          "children": [
            {
              "id": "taxon:pokemon_172_standard",
              "label": ""
            },
            {
              "id": "taxon:pokemon_172_shiny",
              "label": "Shiny"
            }
          ]
        },
        {
          "id": "taxon:pokemon_025",
          "scientificName": "Pikachu",
          "externalReference": [
```

```
        {
          "serviceId": "service:wikidata",
          "externalId": "Q9351"
        },
        {
          "serviceId": "service:example_api",
          "externalId": "Pikachu"
        }
      ],
      "followUp": "https://example.com/pikachu_costumes",
      "children": [
        {
          "id": "taxon:pokemon_025_standard",
          "isEndPoint": true,
          "label": "",
          "children": [
            {
              "id": "taxon:pokemon_025_standard_male",
              "label": "♂"
            },
            {
              "id": "taxon:pokemon_025_standard_female",
              "label": "♀"
            }
          ]
        },
        {
          "id": "taxon:pokemon_025_shiny",
          "label": "Shiny"
        }
      ]
    },
    {
      "id": "taxon:pokemon_026",
      "scientificName": "Raichu",
      "children": [
        {
          "id": "taxon:pokemon_026_standard",
          "label": ""
        },
        {
          "id": "taxon:pokemon_026_shiny",
          "label": "Shiny"
        }
      ]
    }
  ]
},
{
  "id": "taxon:pokemon_115",
  "scientificName": "Kangaskhan",
  "geography": {
```

```
    "polygon": [
      [
        [
          [
            130.0341796875,
            -10.228437266155943
          ],
          [
            111.97265625,
            -21.779905342529634
          ],
          [
            115.09277343749999,
            -36.91476428895593
          ],
          [
            131.2646484375,
            -32.916485347314385
          ],
          [
            141.8994140625,
            -40.44694705960048
          ],
          [
            150.82031249999997,
            -38.8225909761771
          ],
          [
            154.95117187499997,
            -26.15543796871355
          ],
          [
            142.3388671875,
            -10.09867012060338
          ],
          [
            138.69140625,
            -12.382928338487396
          ],
          [
            130.0341796875,
            -10.228437266155943
          ]
        ]
      ]
    ]
  }
},
{
  "id": "taxon:castform",
  "scientificName": "Castform"
}
```

```
],
"characters": [
  {
    "id": "character:type_of_pokemon",
    "title": "Pokémon type",
    "states": [
      {
        "id": "state:pokemon_type_electric",
        "title": "Electric"
      }
    ]
  },
  {
    "id": "character:color",
    "title": "Color of body",
    "states": [
      {
        "id": "state:color_blue",
        "title": "Blue"
      },
      {
        "id": "state:color_red",
        "title": "Red"
      }
    ]
  },
  {
    "id": "character:all_colors",
    "title": "Colors on body of the Pokémon",
    "type": "non-exclusive",
    "states": [
      {
        "id": "state:colors_yellow",
        "title": "Yellow"
      },
      {
        "id": "state:colors_red",
        "title": "Red"
      },
      {
        "id": "state:colors_black",
        "title": "Black"
      },
      {
        "id": "state:colors_brown",
        "title": "Brown"
      },
      {
        "id": "state:colors_white",
        "title": "White"
      }
    ]
```

```
    },
    {
      "id": "character:tail_shape",
      "title": "Shape of the tail end",
      "states": [
        {
          "id": "state:pointy_tail",
          "title": "Pointy",
          "media": "media:pointy_tail"
        },
        {
          "id": "state:lobed_tail",
          "title": "Double-lobed",
          "media": "media:lobed_tail"
        }
      ]
    },
    {
      "id": "character:wearing_hat",
      "title": "Is the Pokémon wearing a hat?",
      "states": [
        {
          "id": "state:hat",
          "title": "Yes"
        },
        {
          "id": "state:no_hat",
          "title": "No"
        }
      ]
    },
    {
      "id": "character:hat_shape",
      "title": "What is the style of the hat?",
      "logicalPremise": "state:hat",
      "states": [
        {
          "id": "state:bowler_hat",
          "title": "Bowler hat"
        },
        {
          "id": "state:top_hat",
          "title": "Top hat"
        }
      ]
    },
    {
      "id": "character:weight",
      "title": "How much does the Pokémon weigh?",
      "userRequirement": "requirement:catch",
      "type": "numerical",
      "min": 1,
```

```
    "max": 125,
    "stepSize": 1,
    "unit": "kg"
  }
],
"statements": [
  {
    "id": "statement:pikachu_is_electric",
    "taxon": "taxon:pokemon_025",
    "character": "character:type_of_pokemon",
    "value": "state:pokemon_type_electric",
    "frequency": 1
  },
  {
    "id": "statement:pikachu_is_never_blue",
    "taxon": "taxon:pokemon_025",
    "character": "character:color",
    "value": "state:color_blue",
    "frequency": 0
  },
  {
    "id": "statement:pikachu_lobed_tail",
    "taxon": "taxon:pokemon_025",
    "character": "character:tail_shape",
    "value": "state:lobed_tail",
    "frequency": 0.5
  },
  {
    "id": "statement:pikachu_pointy_tail",
    "taxon": "taxon:pokemon_025",
    "character": "character:tail_shape",
    "value": "state:pointy_tail",
    "frequency": 0.5
  },
  {
    "id": "statement:pikachu_weight",
    "taxon": "taxon:pokemon_025",
    "character": "character:weight",
    "value": [
      2.98,
      10.1
    ],
    "frequency": 1
  },
  {
    "id": "statement:castform_rain_type",
    "taxon": "taxon:castform",
    "character": "character:castform_type",
    "value": "state:rainy_castform",
    "frequency": 0.2
  },
  {
```

```
  "id": "statement:castform_rain_type_bergen",
  "taxon": "taxon:castform",
  "character": "character:castform_type",
  "value": "state:rainy_castform",
  "frequency": 0.9,
  "geography": {
    "polygon": [
      [
        [
          [
            5.27618408203125,
            60.44976847885747
          ],
          [
            5.218505859375,
            60.4233434866285
          ],
          [
            5.27618408203125,
            60.36160157353732
          ],
          [
            5.395660400390625,
            60.36839212633114
          ],
          [
            5.4052734375,
            60.421309904895715
          ],
          [
            5.27618408203125,
            60.44976847885747
          ]
        ]
      ]
    ]
  }
},
{
  "id": "statement:pikachu_contains_yellow",
  "taxon": "taxon:pokemon_025",
  "character": "character:all_colors",
  "value": "state:colors_yellow",
  "frequency": 1
},
{
  "id": "statement:pikachu_contains_black",
  "taxon": "taxon:pokemon_025",
  "character": "character:all_colors",
  "value": "state:colors_black",
  "frequency": 1
},
```

```
{
  "id": "statement:pikachu_contains_red",
  "taxon": "taxon:pokemon_025",
  "character": "character:all_colors",
  "value": "state:colors_red",
  "frequency": 1
},
{
  "id": "statement:pikachu_contains_no_brown",
  "taxon": "taxon:pokemon_025",
  "character": "character:all_colors",
  "value": "state:colors_brown",
  "frequency": 0
},
{
  "id": "statement:pikachu_contains_no_white",
  "taxon": "taxon:pokemon_025",
  "character": "character:all_colors",
  "value": "state:colors_white",
  "frequency": 0
},
{
  "id": "statement:raichu_contains_yellow",
  "taxon": "taxon:pokemon_026",
  "character": "character:all_colors",
  "value": "state:colors_yellow",
  "frequency": 1
},
{
  "id": "statement:raichu_contains_no_black",
  "taxon": "taxon:pokemon_026",
  "character": "character:all_colors",
  "value": "state:colors_black",
  "frequency": 0
},
{
  "id": "statement:raichu_contains_no_red",
  "taxon": "taxon:pokemon_026",
  "character": "character:all_colors",
  "value": "state:colors_red",
  "frequency": 0
},
{
  "id": "statement:raichu_contain_brown",
  "taxon": "taxon:pokemon_026",
  "character": "character:all_colors",
  "value": "state:colors_brown",
  "frequency": 1
},
{
  "id": "statement:raichu_contains_white",
  "taxon": "taxon:pokemon_026",
```

```
        "character": "character:all_colors",
        "value": "state:colors_white",
        "frequency": 1
      }
    ],
    "userRequirements": [
      {
        "id": "requirement:catch",
        "title": "Catching required",
        "warning": "To answer this, you have to catch the Pokémon
first.",
        "description": "1. Select a pokéball color.\n2. Hold the ball,
spinning it a few times.\n3. Fling the pokéball towards the
Pokémon, adjusting for the curveball generated from spinning the
ball.\n4. Try to hit the circle when it is at its smallest.",
        "descriptionUrl":
"https://niantic.helpshift.com/hc/en/6-pokemon-go/faq/102-finding-catc
hing-wild-pokemon/"
      }
    ],
    "externalServices": [
      {
        "id": "service:wikidata",
        "title": "Wikidata",
        "url": "https://www.wikidata.org/w/api.php"
      },
      {
        "id": "service:example_api",
        "title": "Example",
        "description": "Gives the probability for a taxon, given its
weight and location.",
        "url": "https://api.example.com"
      }
    ],
    "mediaElements": [
      {
        "id": "media:pointy_tail",
        "mediaElement": {
          "file": [
            {
              "url":
"https://github.com/WouterKoch/Clavis/raw/main/Keys/Images/pointy_100.
png",
              "width": 100,
              "height": 100,
              "license":
"https://creativecommons.org/licenses/by/4.0/",
              "creator": "person:wouterkoch"
            },
            {
              "url":
"https://github.com/WouterKoch/Clavis/raw/main/Keys/Images/pointy_250.
```

```
png",
            "width": 250,
            "height": 250,
            "license":
"https://creativecommons.org/licenses/by/4.0/",
            "creator": "person:wouterkoch"
          }
        ]
      }
    },
    {
      "id": "media:lobed_tail",
      "mediaElement": {
        "file": [
          {
            "url":
"https://github.com/WouterKoch/Clavis/raw/main/Keys/Images/lobed_100.p
ng",
            "width": 100,
            "height": 100,
            "license":
"https://creativecommons.org/licenses/by/4.0/",
            "creator": "person:wouterkoch"
          },
          {
            "url":
"https://github.com/WouterKoch/Clavis/raw/main/Keys/Images/lobed_250.p
ng",
            "width": 250,
            "height": 250,
            "license":
"https://creativecommons.org/licenses/by/4.0/",
            "creator": "person:wouterkoch"
          }
        ]
      }
    }
  ]
}
```
