## Supporting information File 3. Clavis key example: titmice for "Clavis: an open and versatile identification key format"


```
{
   "$schema":
"https://raw.githubusercontent.com/Artsdatabanken/Clavis/main/Schema/C
lavis.json",
   "title": "A key to titmice in Norway",
   "language": "en",
   "license": "https://creativecommons.org/licenses/by/4.0/",
   "creator": "person:wouterkoch",
   "lastModified": "2022-03-19 21:41:00",
   "identifier": "9be11d7e-c147-400a-899e-b3d5e4bcc6a1",
   "geography": {
      "name": "Norway",
      "polygon": [
         [
            [
               [
                  33.22265625,
                  69.56522590149099
               ],
               [
                  29.267578125,
                  71.15939141681443
               ],
               [
                  23.5546875,
                  71.28669893545877
               ],
               [
                  17.2265625,
                  69.97549253616164
               ],
               [
                  12.392578125,
                  68.366801093914
               ],
               [
                  11.2939453125,
                  65.91062334197893
               ],
               [
                  3.8671874999999996,
                  62.103882522897855
               ],
               [
                  4.7021484375,
                  58.516651799363785
```

```
        ],
        [
          7.119140625,
          57.70414723434193
        ],
        [
          11.953125,
          58.83649009392136
        ],
        [
          13.3154296875,
          61.39671887310411
        ],
        [
          12.8759765625,
          63.6267446447533
        ],
        [
          14.94140625,
          64.07219957867282
        ],
        [
          15.2490234375,
          66.05371622067922
        ],
        [
          18.852539062499996,
          68.02402198693447
        ],
        [
          25.048828125,
          68.51214331858073
        ],
        [
          26.894531249999996,
          69.54987728327795
        ],
        [
          29.003906249999996,
          68.8159271333607
        ],
        [
          33.22265625,
          69.56522590149099
        ]
      ]
    ]
  ]
},
"persons": [
  {
    "id": "person:wouterkoch",
```

```
         "name": "Wouter Koch"
      }
   ],
   "externalServices": [
      {
         "id": "service:nbic_taxa",
         "title": "NBIC taxonomy scientificNameId",
         "description": "To retrieve taxon information based on the
NBIC scientificNameId, e.g. through
https://www.artsdatabanken.no/api/Taxon/ByScientificNameId/4362",
         "provider": "Norwegian Biodiversity Information Centre",
         "url": "https://www.artsdatabanken.no/help"
      }
   ],
   "taxa": [
      {
         "id": "taxon:paridae",
         "scientificName": "Paridae",
         "rank": "family",
         "vernacularName": "titmice",
         "externalReference": {
            "serviceId": "service:nbic_taxa",
            "externalId": "4362"
         },
         "children": [
            {
               "id": "taxon:cyanistes",
               "scientificName": "Cyanistes",
               "rank": "genus",
               "externalReference": {
                  "serviceId": "service:nbic_taxa",
                  "externalId": "4364"
               },
               "children": [
                  {
                     "id": "taxon:cyanistes_caeruleus",
                     "scientificName": "Cyanistes caeruleus",
                     "vernacularName": "blue tit",
                     "rank": "species",
                     "externalReference": {
                        "serviceId": "service:nbic_taxa",
                        "externalId": "4365"
                     }
                  }
               ]
            },
            {
               "id": "taxon:lophophanes",
               "scientificName": "Lophophanes",
               "rank": "genus",
               "externalReference": {
                  "serviceId": "service:nbic_taxa",
```

```
        "externalId": "4368"
      },
      "children": [
        {
          "id": "taxon:lophophanes_cristatus",
          "scientificName": "Lophophanes cristatus",
          "vernacularName": "crested tit",
          "rank": "species",
          "externalReference": {
            "serviceId": "service:nbic_taxa",
            "externalId": "4369"
          },
          "geography": {
            "polygon": [
              [
                [
                  [
                    11.997070312499998,
                    58.74540696858028
                  ],
                  [
                    13.0517578125,
                    59.977005492196
                  ],
                  [
                    12.8759765625,
                    63.31268278043484
                  ],
                  [
                    19.2041015625,
                    68.57644086491786
                  ],
                  [
                    13.4912109375,
                    68.78414378041504
                  ],
                  [
                    9.7119140625,
                    64.47279382008166
                  ],
                  [
                    4.21875,
                    62.2679226294176
                  ],
                  [
                    3.69140625,
                    59.40036514079251
                  ],
                  [
                    6.723632812499999,
                    57.70414723434193
                  ],
```

```
                     [
                        11.997070312499998,
                        58.74540696858028
                     ]
                  ]
               ]
            ]
         }
      }
   ]
},
{
   "id": "taxon:parus",
   "scientificName": "Parus",
   "rank": "genus",
   "externalReference": {
      "serviceId": "service:nbic_taxa",
      "externalId": "4363"
   },
   "children": [
      {
         "id": "taxon:parus_major",
         "scientificName": "Parus major",
         "vernacularName": "great tit",
         "rank": "species",
         "externalReference": {
            "serviceId": "service:nbic_taxa",
            "externalId": "4372"
         }
      }
   ]
},
{
   "id": "taxon:periparus",
   "scientificName": "Periparus",
   "rank": "genus",
   "externalReference": {
      "serviceId": "service:nbic_taxa",
      "externalId": "4374"
   },
   "children": [
      {
         "id": "taxon:periparus_ater",
         "scientificName": "Periparus ater",
         "vernacularName": "coal tit",
         "rank": "species",
         "externalReference": {
            "serviceId": "service:nbic_taxa",
            "externalId": "4375"
         }
      }
   ]
```

```
},
{
   "id": "taxon:poecile",
   "scientificName": "Poecile",
   "rank": "genus",
   "externalReference": {
      "serviceId": "service:nbic_taxa",
      "externalId": "4378"
   },
   "children": [
      {
         "id": "taxon:poecile_palustris",
         "scientificName": "Poecile palustris",
         "vernacularName": "marsh tit",
         "rank": "species",
         "externalReference": {
            "serviceId": "service:nbic_taxa",
            "externalId": "4385"
         },
         "geography": {
            "polygon": [
               [
                  [
                     [
                        11.997070312499998,
                        58.74540696858028
                     ],
                     [
                        13.0517578125,
                        59.977005492196
                     ],
                     [
                        12.8759765625,
                        63.31268278043484
                     ],
                     [
                        19.2041015625,
                        68.57644086491786
                     ],
                     [
                        13.4912109375,
                        68.78414378041504
                     ],
                     [
                        9.7119140625,
                        64.47279382008166
                     ],
                     [
                        4.21875,
                        62.2679226294176
                     ],
                     [
```

```
                      3.69140625,
                      59.40036514079251
                    ],
                    [
                      6.723632812499999,
                      57.70414723434193
                    ],
                    [
                      11.997070312499998,
                      58.74540696858028
                    ]
                  ]
                ]
              ]
            }
          },
          {
            "id": "taxon:poecile_montanus",
            "scientificName": "Poecile montanus",
            "vernacularName": "willow tit",
            "rank": "species",
            "externalReference": {
              "serviceId": "service:nbic_taxa",
              "externalId": "4382"
            }
          },
          {
            "id": "taxon:poecile_cinctus",
            "scientificName": "Poecile cinctus",
            "vernacularName": "Siberian tit",
            "rank": "species",
            "externalReference": {
              "serviceId": "service:nbic_taxa",
              "externalId": "4379"
            }
          }
        ]
      }
    ]
  }
],
"characters": [
  {
    "id": "character:head_top",
    "title": "Top of the head",
    "states": [
      {
        "id": "state:black_or_dark_grey",
        "title": "Black or dark grey"
      },
      {
        "id": "state:blue",
```

```
      "title": "Blue"
   },
   {
      "id": "state:speckled_crest",
      "title": "Speckled black and white, with a crest"
   }
  ]
},
{
  "id": "character:head_top",
  "title": "Top of the head",
  "states": [
   {
      "id": "state:black_or_dark_grey",
      "title": "Black or dark grey"
   },
   {
      "id": "state:brown",
      "title": "Brown"
   },
   {
      "id": "state:blue",
      "title": "Blue"
   },
   {
      "id": "state:speckled_crest",
      "title": "Speckled black and white, with a crest"
   }
  ]
},
{
  "id": "character:chest_color",
  "title": "Color of the chest",
  "states": [
   {
      "id": "state:yellow",
      "title": "Yellow"
   },
   {
      "id": "state:grey_brown",
      "title": "Grey to brown"
   }
  ]
},
{
  "id": "character:wing_bar",
  "title": "White bar on the wing",
  "states": [
   {
      "id": "state:wing_bar",
      "title": "Present"
   },
```

```
        {
          "id": "state:no_wing_bar",
          "title": "Absent"
        }
      ]
    },
    {
      "id": "character:black_of_cheek",
      "title": "Color of the cheek at the back",
      "states": [
        {
          "id": "state:white_cheek_back",
          "title": "Entire cheek white"
        },
        {
          "id": "state:brown_cheek_back",
          "title": "Sullied brown"
        }
      ]
    },
    {
      "id": "character:wing_secondaries_color",
      "title": "Color of secondary wing feathers",
      "states": [
        {
          "id": "state:secondaries_pale",
          "title": "Paler than rest of the wing"
        },
        {
          "id": "state:secondaries_not_pale",
          "title": "No clear different from rest of the wing"
        }
      ]
    }
  ],
  "statements": [
    {
      "id": "statement:crested_tit_head",
      "taxon": "taxon:lophophanes_cristatus",
      "character": "character:head_top",
      "value": "state:speckled_crest",
      "frequency": 1
    },
    {
      "id": "statement:blue_tit_head",
      "taxon": "taxon:cyanistes_caeruleus",
      "character": "character:head_top",
      "value": "state:blue",
      "frequency": 1
    },
    {
      "id": "statement:poecile_palustris_head",
```

```
        "taxon": "taxon:poecile_palustris",
        "character": "character:head_top",
        "value": "state:black_or_dark_grey",
        "frequency": 1
    },
    {
        "id": "statement:poecile_montanus_head",
        "taxon": "taxon:poecile_montanus",
        "character": "character:head_top",
        "value": "state:black_or_dark_grey",
        "frequency": 1
    },
    {
        "id": "statement:poecile_cinctus_head",
        "taxon": "taxon:poecile_cinctus",
        "character": "character:head_top",
        "value": "state:brown",
        "frequency": 1
    },
    {
        "id": "statement:great_tit_head",
        "taxon": "taxon:parus_major",
        "character": "character:head_top",
        "value": "state:black_or_dark_grey",
        "frequency": 1
    },
    {
        "id": "statement:coal_tit_head",
        "taxon": "taxon:periparus_ater",
        "character": "character:head_top",
        "value": "state:black_or_dark_grey",
        "frequency": 1
    },
    {
        "id": "statement:crested_tit_chest",
        "taxon": "taxon:lophophanes_cristatus",
        "character": "character:chest_color",
        "value": "state:grey_brown",
        "frequency": 1
    },
    {
        "id": "statement:blue_tit_chest",
        "taxon": "taxon:cyanistes_caeruleus",
        "character": "character:chest_color",
        "value": "state:yellow",
        "frequency": 1
    },
    {
        "id": "statement:poecile_chest",
        "taxon": "taxon:poecile",
        "character": "character:chest_color",
        "value": "state:grey_brown",
```

```
    "frequency": 1
},
{
    "id": "statement:great_tit_chest",
    "taxon": "taxon:parus_major",
    "character": "character:chest_color",
    "value": "state:yellow",
    "frequency": 1
},
{
    "id": "statement:coal_tit_chest",
    "taxon": "taxon:periparus_ater",
    "character": "character:chest_color",
    "value": "state:grey_brown",
    "frequency": 1
},
{
    "id": "statement:crested_tit_bar",
    "taxon": "taxon:lophophanes_cristatus",
    "character": "character:wing_bar",
    "value": "state:no_wing_bar",
    "frequency": 1
},
{
    "id": "statement:blue_tit_bar",
    "taxon": "taxon:cyanistes_caeruleus",
    "character": "character:wing_bar",
    "value": "state:wing_bar",
    "frequency": 1
},
{
    "id": "statement:poecile_bar",
    "taxon": "taxon:poecile",
    "character": "character:wing_bar",
    "value": "state:no_wing_bar",
    "frequency": 1
},
{
    "id": "statement:great_tit_bar",
    "taxon": "taxon:parus_major",
    "character": "character:wing_bar",
    "value": "state:wing_bar",
    "frequency": 1
},
{
    "id": "statement:coal_tit_bar",
    "taxon": "taxon:periparus_ater",
    "character": "character:wing_bar",
    "value": "state:wing_bar",
    "frequency": 1
},
{
```

```
            "id": "statement:poecile_palustris_cheek_back",
            "taxon": "taxon:poecile_palustris",
            "character": "character:black_of_cheek",
            "value": "state:brown_cheek_back",
            "frequency": 1
         },
         {
            "id": "statement:poecile_montanus_cheek_back",
            "taxon": "taxon:poecile_montanus",
            "character": "character:black_of_cheek",
            "value": "state:white_cheek_back",
            "frequency": 1
         },
         {
            "id": "statement:poecile_palustris_wing_secondaries_color",
            "taxon": "taxon:poecile_palustris",
            "character": "character:wing_secondaries_color",
            "value": "state:secondaries_not_pale",
            "frequency": 1
         },
         {
            "id": "statement:poecile_montanus_wing_secondaries_color",
            "taxon": "taxon:poecile_montanus",
            "character": "character:wing_secondaries_color",
            "value": "state:secondaries_pale",
            "frequency": 1
         }
      ]
   }
```
